# Capillary-induced bundling of biopolymer networks via protein condensates

**DOI:** 10.1101/2025.01.22.634359

**Authors:** Carolyn A. Feigeles, Artis Brasovs, Adam Puchalski, Olivia Laukat, Konstantin G. Kornev, Kimberly L. Weirich

## Abstract

Within the cell, biopolymers form self-organized assemblies that regulate cellular processes. These assemblies can be constructed through either protein interactions or phase separation. It is known that actin filaments, which form the mechanical structure of cells, are assembled into networks and bundles by protein cross-linkers. Different network and bundle microstructures support different physiological functions. Recently, there is evidence that protein condensates interact with biopolymers to create bundles. Here, we show that protein condensates colocalize with actin filaments and form networks of bundles. The condensates absorb on actin bundles and relax into a barrel shaped droplet on bundles, evocative of drops of simple liquids on fibers. We investigate the condensate spreading and measure contact angle that condensates make with bundles. Condensates at the intersection of bundles cause capillary bridges which induce network remodeling. Our results suggest that network formation, bundling, and remodeling in biopolymer assemblies could be induced by capillary interactions due to condensates. Understanding this bundling mechanism could expand our toolkit for making self-assembled fiber-based soft materials.

## Introduction

An important class of biological materials are the dynamic protein assemblies in cells, which regulate mechanical processes, such as extending protrusions and controlling shape, exerting forces, and cargo transport (1, 2). These vastly different physiological processes are mediated by a small number of cytoskeletal proteins that organize into a myriad of microstructures (3, 4). Actin filaments arrange into networks, bundles, and networks of bundles. It is well known that protein cross-linkers, which have two or more domains that bind to actin, arrange actin into the various assemblies, where the microstructure derives from the cross-linker properties.

There are many other protein assemblies in the cytoplasm, such as liquid-liquid phase separated condensates (5–7). The liquid-like behavior of protein condensates suggest possible interactions between different surfaces within the cytoplasm, such as wetting vesicles and filaments. It has recently been suggested that protein condensates could wet cellular assemblies, such as vesicles and filaments, and induce higher-order structure formation via capillary-like interactions (8–10). When the bending energy of fibers is small compared to the interfacial energy, the fiber could wrap around the drop or be engulfed inside it (11). When the bending energy is large, the drop could bead or spread on the fiber. When multiple wettable fibers interact with the same drop, capillary bridges form between them, drawing the fibers together (12). Despite extensive research on drops interacting with fiber that have a cross-section much greater than the scale of the liquid molecules (13–15), little is known about how drops interact with materials that have length scale on the order of the liquid molecules, such as protein drops interacting with protein-based materials.

Here, we investigate the interaction of protein condensates with actin filaments. We find that actin filaments and condensates colocalize into bundles that bridge together into a space spanning network. By observing the interaction of a single condensate and a bundle of actin filaments, we find that condensate droplets spread along a bundle evocative of a macroscopic unduloidal drop on fiber (13). Our results suggest a capillary-based mechanism for cytoskeletal assembly formation, distinct from cross-linker induced bundling.

## Results

### Condensates and filaments colocalize in bundles

To create actin networks, we polymerize actin filaments by adding 2.64 μM actin monomer to actin polymerization buffer in the presence of a depletion agent, which causes filaments to accumulate at the surfaces of the sample chamber (Fig. 1A and 1B) (16). To the pre-polymerized actin, we add 4 μM FUS protein. At this concentration, FUS without actin forms protein condensates in buffer conditions (Fig. 1C), consistent with liquid-liquid phase-separated condensates reported previously for similar conditions (17). Upon adding FUS condensates to the actin network, actin filaments rapidly begin to bundle (Fig. 1D). First, small segments of condensate (gray, and blue in composite) accumulate between actin filaments, inducing bundling (0-40 s). Eventually, the condensates cover the length of the bundles (>200 s). Inspecting the intensity along a line perpendicular to the bundle length reveals that condensates and actin have maxima at the same location (Fig. 1E). This indicates that the two proteins are colocalized in bundles, suggesting that actin filaments coexist with condensates in liquid columns. The average bundle thickness is 0.23 ± 0.10 μm (See SI). This is comparable to the average thickness of actin bundles formed by physiological cross-linkers (18, 19).

**Figure 1:**
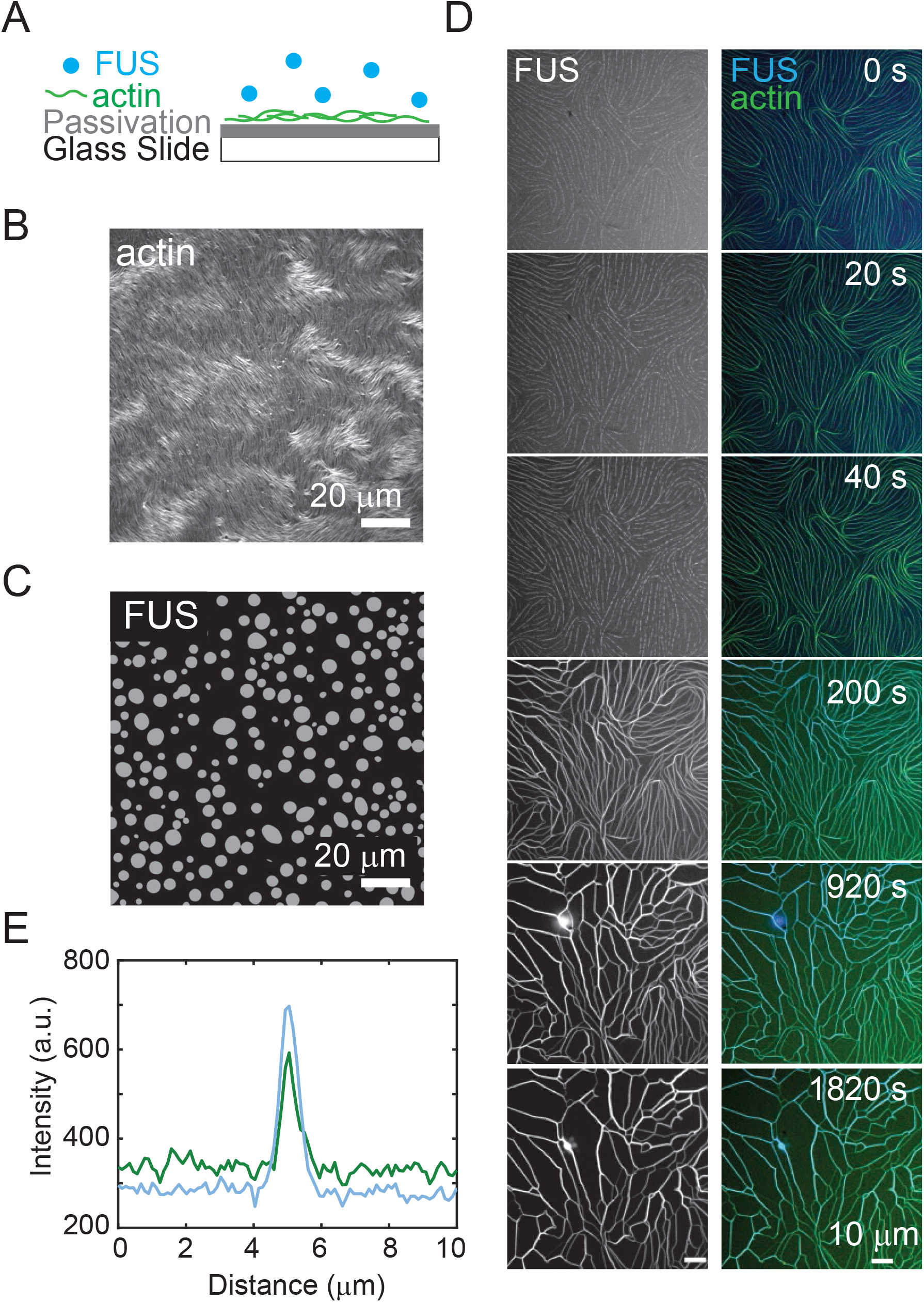
FUS condensates and actin filaments colocalize in bundles. (A) Schematic of experimental setup. FUS condensates (blue) are added to actin filaments (green) crowded to a passivated surface. (B) Fluorescence microscopy image of entangled actin network (2.64 µM) crowded to a surface. Scale bar is 20 μm. (C) Fluorescence microscopy image of FUS condensates (4 µM). Scale bar is 20 μm. (D) Fluorescence images of FUS condensates (4 µM FUS) and actin network bundling and remodeling over time. Left column images are FUS (grey). Right column is FUS (blue) and actin (green) merged. Scale bar is 10 μm. (E) Distribution of intensity of FUS and actin over a given distance, showing colocalization within the network.

### Condensates adsorb on bundles and form drops

When an individual condenstate in solution approaches actin previously bundled by condensate, the condensate spreads on the bundle (Fig. 2A). The free condensate is initially spherical before it absorbs to the bundle (Fig. 2A, 0-15 s), and changes morphology as it spreads into a final, barrel-shape (15-60 s). To characterize the condensate spreading, we measure the aspect ratio, 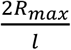, of the condensate over time, where *R*_*max*_ and *l* are defined in Fig. 2B. The aspect ratio is initially ∼1 prior to the condensate adsorbing onto the actin bundle, consistent with a spherical droplet. As the condensate spreads on the bundle, the aspect ratio decreases to ∼0.7, with a characteristic decay time of ∼5.8 s from an exponential fit. The condensate remains barrel-shaped on the bundle until the end of observation (∼200 s), suggesting that the bundle cannot accomodate more condensate incorporation after the condensate has spread to a steady state morphology.

**Figure 2:**
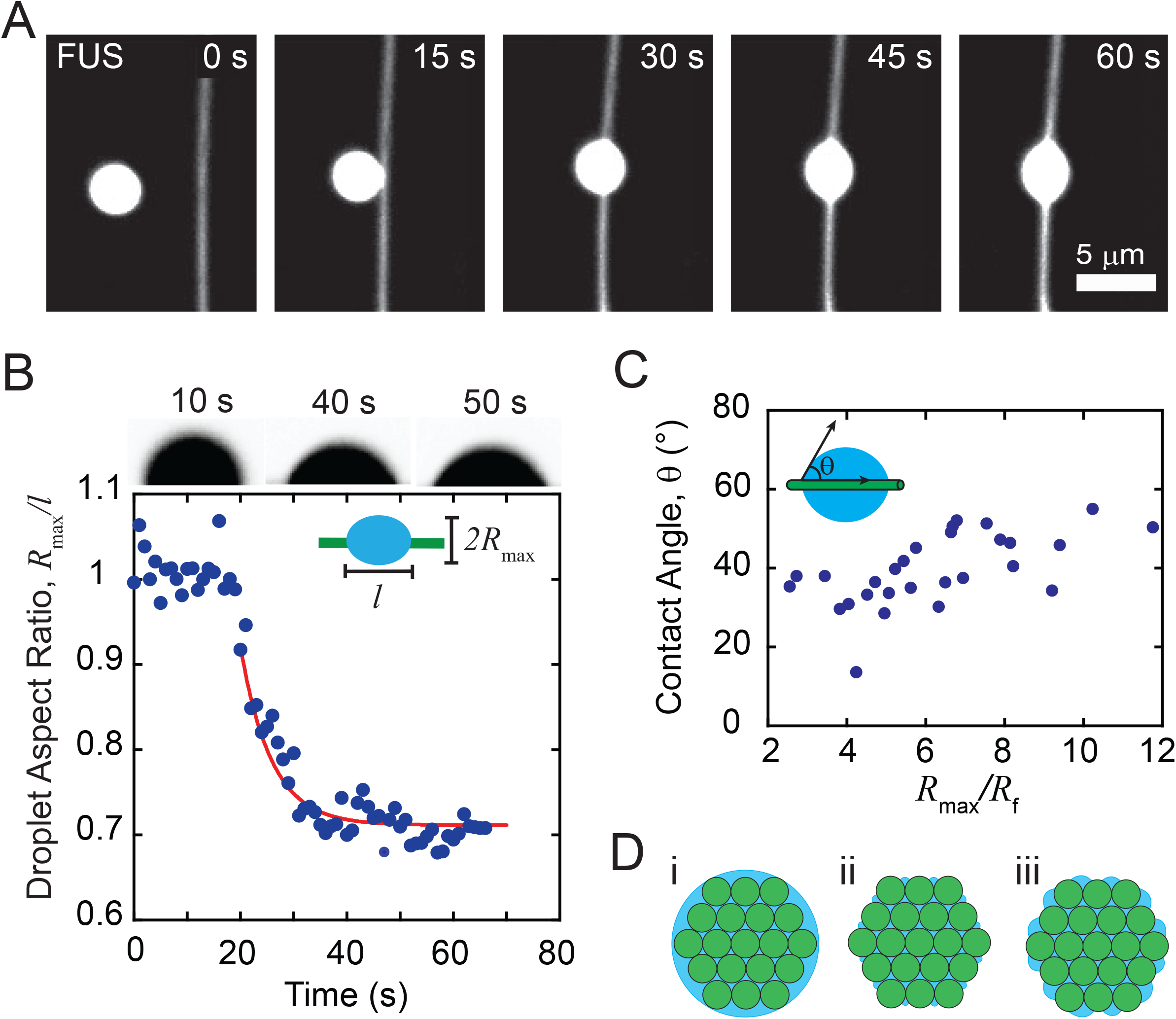
FUS condensates adsorb on actin bundles and form barrel-shaped condensates. (A) Fluorescence microscopy images of a FUS condensate absorbing to and spreading over an actin bundle. The final sample concentration is 2.64 μM actin and 4.4 μM FUS. Scale bar is 5 μm. (B) Aspect ratio (2*R*_max_/*l*) of condensate in (A) as a function of time. Images of condensate cropped at the bundle interface over time (top). (C) Schematic of contact angle definition, θ (inset). Contact angle as a function of the condensate radius, *R*_max_, rescaled by the actin bundle radius, *R*_f_ (n=28 condensates). (D) Cartoon of different potential condensate-filament surfaces in bundles. i) the filaments are bundled inside a column of condensate. ii) filaments and condensates coexist at the surface, forming a surface where filaments can be directly in contact with buffer or condensates form liquid bridges that meet actin filaments at acute contact angles. iii) filaments and condensates coexist at the surface as in (ii), but liquid bridges have larger radius of curvature.

We measure the contact angle between the condensate and the bundle, which provides information regarding relative surface energies of components (20, 21). To measure the contact angle (Fig. 2C, inset), we first extract the condensate contour and fit to an unduloid (Fig. S1). Across 28 condensates, the contact angle is scattered without dependence on condensate radius (Fig. S2). Normalizing the condensate radius, *R*_*max*_, by the fiber radius, *R*_*f*_, reveals that the contact angle remains constant across a range of condensate sizes with respect to bundle radius (Fig. 2C, n=28). The low average contact angle, *θ* = 39.5° ± 9.01°, indicates in these buffer conditions, that it is favorable for FUS to wet bundles (Fig. 2A, (22)). A contact angle closer to zero might have been expected, in the limit where condensates completely coat the bundle, since the condensates are spreading on bundles that condensates have already wet (Fig. 2D i). The deviation of the contact angle from zero suggests that some parts of the actin within the bundle are directly exposed to the buffer.

To distinguish between cases where the condensates are completely or partially coating the bundle (Fig. 2D), we consider the capillary pressure which is the pressure between two immiscible fluids at at interface (8, 14, 23). The capillary presssure, Δ*P*_condensate_, between the condensate and bundle for an unduloid drop is 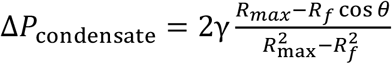, where γ is the interfacial tension of condensates (13). Assuming an interfacial tension for FUS condensates of 90 μNm^-1^ (24), we found that the capillary pressure ranges from ∼20 to 70 Nm^-2^ (Fig. S4). In the case where the condensate completely wets the bundle, Δ*P*_condensate_, would be expected to be equal to the capillary pressure of cylindrical FUS column, Δ*P*_column_ = γ/*RR*_f_. We find that pressure range in the condensate is smaller than the pressure range associated with a FUS column, ∼65 to 260 Nm^-2^. Plotting the dimensionless capillary pressure, 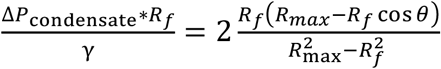, which is the capillary pressure normalized by the capillary pressure of the cylindrical FUS column, against the condensate radius rescaled by the fiber radius,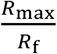, reveals that for majority of the tested droplets, this ratio is less than 1 (Fig. S5), indicating that the completely wet case shown in Fig. 2Di is inconsistent with the data. In contrast, previous studies of wetting phenomena on yarns made of wettable nanofibers showed complete drop absorption (25–28).

To better understand how a barrel-shaped condensate with a positive capillary pressure could be in equilibrium with a bundle of wettable nanofibers that is not completely covered by the fluid column, we consider another scenario of FUS absorption where each condensate meniscus bridging two actin filaments makes an arc of radius *R*_*FUS*_ (Fig. 2D ii and iii). This radius satisfies the equilibrium condition 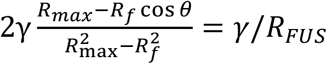. The data *RR*_*max*_/*R*_*f*_ in Fig. 2C, indicates that *R*_*FUS*_ ranges between 2.3 *R*_*f*_ < *R*_FUS_ < 5.4*R*_*f*_, suggesting that menisci in the surface grooves of the bundle could have a greater radius of curvature while in equilibrium with the droplet. To evaluate the contact angle that the FUS makes with individual actin filament, the Cassie-Baxter theory designed for this case requires the knowledge of the filament density in the bundle (27, 29), which is not available with the current data.

### Condensates coalesce on bundles

In addition to condensates adsorbing on bundles and spreading, condensates also come in contact with another condensate that has previously spread on an actin bundle (Fig. 3A). When a spherical condensate approaches a barrel-shaped condensate on a bundle, both condensates quickly (<5 s) merge and transiently form an asymmetric condensate on the bundle (40-45 s). The condensate then redistributes, becoming more symmetric about the bundle as it spreads, finally adopting a barrel shape similar to the adsorbed condenstate before coalescence (50 s).

**Figure 3:**
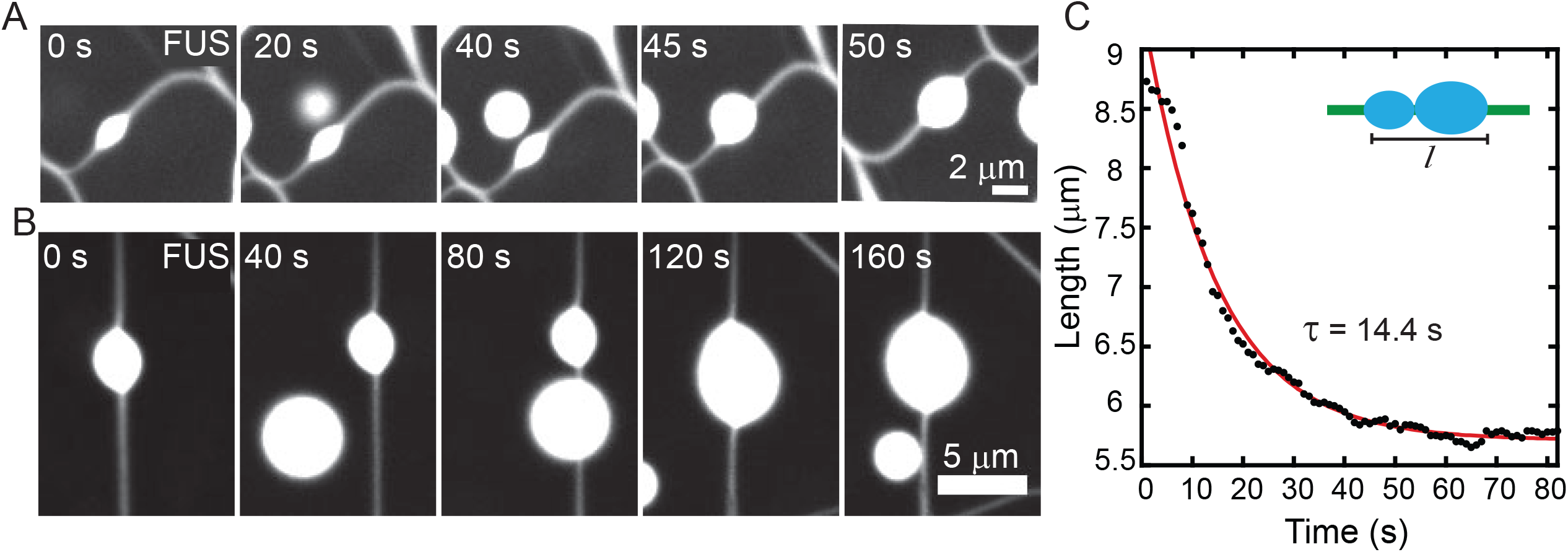
FUS Condensates Coalesce on Filaments and Bundles. (A) Fluorescence microscopy images of a FUS condensate free in solution merging with a barrel-shaped FUS condensate on bundle (2.64 μM actin and 5.28 μM FUS). Scale bar is 2 μm. (B) Fluorescence microscopy images of a FUS condensate spreading on a bundle and merging with a previously absorbed FUS condensate (2.64 μM actin and 4.4 μM FUS). Scale bar is 5 μm. (C) The length of two condensates coalescing on an actin bundle (data in B) as a function of time (black filled circles). Exponential fit (red solid line) has a characteristic decay of τ∼14.4 s.

Another case is when two condensates, which have separately adsorbed and spread on a bundle, coalesce. In Figure 3B, a bundle initially (0 s) has a condensate spread into a barrel shape. When a second larger condensate comes into the focal plane and adsorbs onto the bundle a few microns away from the first (Fig. 3B, 40-120 s), we see it spread. Despite the smaller condensate having a greater capillary pressure than the larger condensate, it does not incorporate into the bundle suggesting that the bundle is fully saturated with the FUS condensate. As the larger condensate spreads, the smaller condensate merges and relaxes into a single barrel-shaped condensate with the sum of the volumes (Fig. 3B, 160 s). We measure the condensate length along the major axis as the two condensates coalesce. The condensate length shortening is consistent with exponential decay that has relaxation time, τ =14.4 s (Fig. 3C). These data indicate condensates are able to coalesce on bundles similarly to how condensates coalesce with each other in solution. The condensate on the bundle maintains properties of a liquid, as would be expected of a drop of a simple liquid on a fiber.

### Condensates bundle filaments

Condensates not only absorb on isolated bundles, but can also exist at the intersection of multiple actin filaments or bundles. In Figure 4A, condensates at the intersection of bundles (0 min), spread on individual bundles and begin to merge (3-6 min). As they coalesce (Fig. 4A, 9-13 min), the actin bundles and filaments are bridged by the condensate and drawn closer together, until they merge into a larger bundle. This indicates that capillary bridges induce bundling in and remodeling of networks of actin filaments bundled by condensates (Fig. 4B).

**Figure 4:**
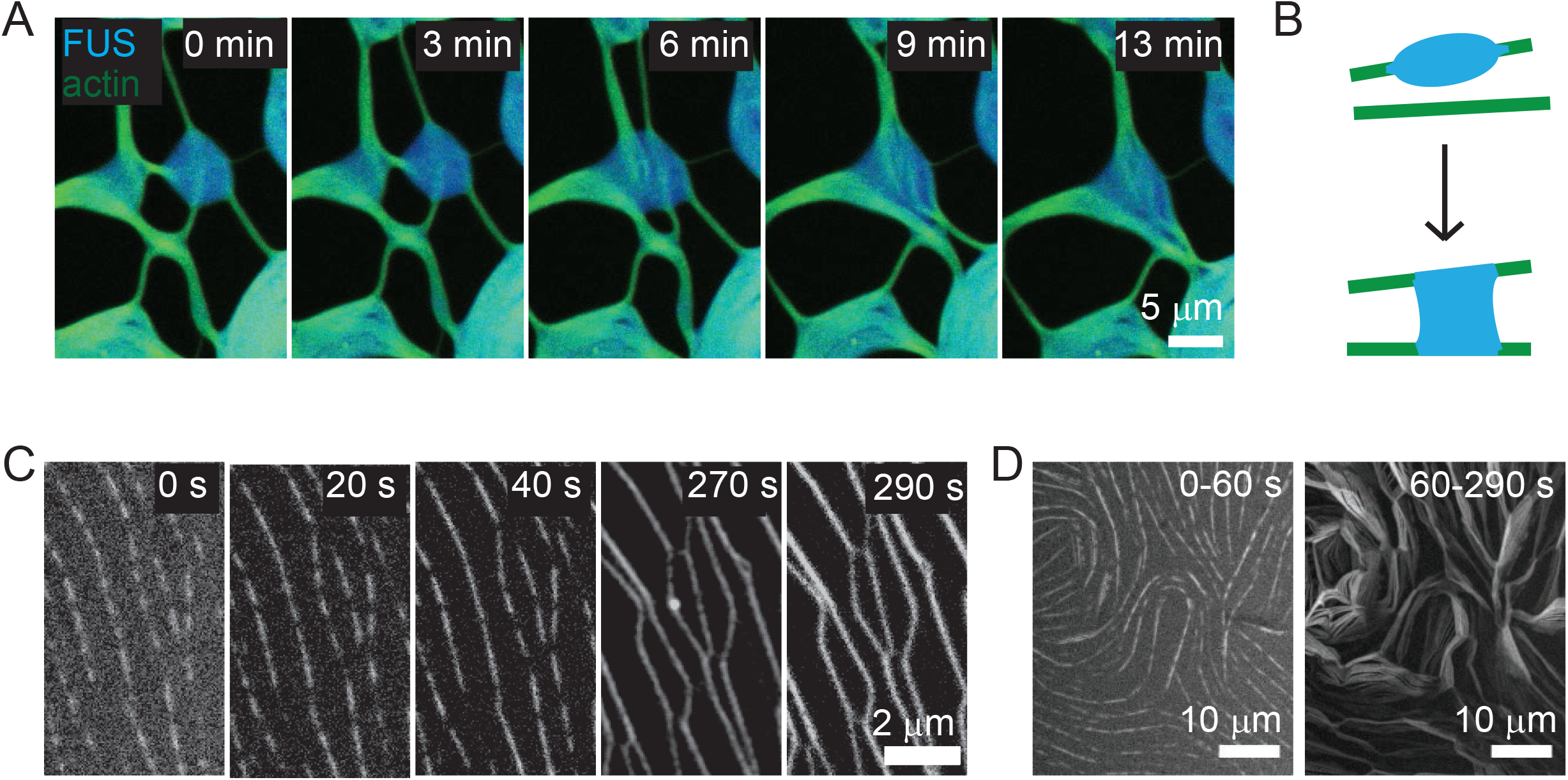
FUS condensates bundle and remodel actin networks via capillary bridges. (A) Fluorescence microscopy images of FUS condensates (blue) at intersection of multiple actin (green) filaments bundled by condensate (0 min). Condensates spread and begin to coalesce (3-6 min), bringing filaments together (9-13 min). Sample concentration is 2.64 μM actin and 2.64 μM FUS. Scale bar is 5 μm. (B) Cartoon of capillary bridging of a condensate (blue) between actin filaments (green). (C) Fluorescence microscopy images of absorbed FUS condensates spreading along and bundling actin filaments. Data is magnified region of data in Figure 1D. Scale bar is 2 μm. (D) Maximum intensity projection of the network shown in Fig. 4B. The initial projection (0-60 s) shows little change in the network structure, while the later frame projection (60-290 s) shows significant network coarsening.

The capillarity-induced bundling observed in networks provides insight into mechanism of initial actin filament bundling by condensates. We have not captured condenstates interacting with single actin filaments, due to the quick formation of initial bundles. However, we do have evidence that condensates can accumulate in the initial stages of actin bundles in small regions (Fig. 4C, a zoomed in region of data in Fig. 1D), reminiscent of capillary bridge segments formed by droplets between two fibers (12). On the order of tens of seconds, these segments become longer while condensates continue to accumulate in the bundles. A maximal intensity projection of the first 60 s, indicates that the overall network structure is stable during intial bundle formation (Fig. 4D, 0-60 s, compare with Fig. 4C, 0-40 s). However, after condensates have spread throughout actin bundles, the bundles begin merging with neighboring bundles and the network coarsens (Fig. 4C, > 200 s). A maximal intensity projection of data from 60-290 s shows bundle mobility and large changes in network structure (Fig. 4D, 60-290 s).

## Discussion

Cytoskeletal assemblies enable a variety of critical physiological functions, supported through variations in microstructures, which have mechanical properties that derive from structure. In cytoskeletal assemblies, the microstructure is predominately known to be regulated through proteins which cross-link cytoskeletal filaments, such as actin and microtubules. Earlier research has reported interaction between FUS condensates and actin filaments, where short (∼100 nm) actin filaments were found to incorporate into FUS condensates, imparting an elasticity that influenced the condensate shape (30). It has been recently reported that cytoskeletal filaments nucleated within protein condensates result in elongated bundles of filaments and condensate such as microtubules with tau condenstates, and actin with and VASP and abLIM1 condensates (31, 17, 32, 33). Actin also forms bundled rings inside of VASP condensates, similar to reports of actin in polyelectrolyte coacervates (17, 32, 34). It has also been reported that condensates and filaments will form larger assemblies, such as bundled networks with abLIM1 and aster shaped bundle assemblies with VASP (33, 32). We present evidence that actin can be bundled by condensates via capillary interactions, similar to drops on fibers, and form networks which remodel through capillary bridges and condenstate coalescence. Capillary bridging and condenstate coalescence are physical mechanisms of network remodeling that are consistent with reports of “zippering” bundles in other condensate-filament systems (17). The remodeling and environmental responsiveness of conventional, cross-linked actin networks derive from cross-linkers ability to dynamically bind and unbind (35). Capillarity-induced bundling and network remodeling may yield networks with not only different microstructure, but dynamics and properties than cross-linked assemblies.

We find that condensates absorb and spread on actin filaments, forming barrel-shaped condensates. Notably, the shape is similar to that characteristic of liquid drops collecting on macroscopic fibers, despite the different length scale of actin filaments (diameter is ∼ 7 nm) and molecular details of protein-based liquids (13, 14). In this research, we have done initial studies with one type of protein condensate (FUS) interacting with the cytoskeletal filament actin. However, there are many types of protein condensates and cytoskeletal filaments, with different chemistries and structures (36–38). Future research might explore the molecular mechanisms of FUS protein interactions with actin, as well as other protein condensates to gain insight into the absorption and spreading process. It would also be interesting to understand if polymer complex coacervates (34, 39–41), which can exhibit liquid behavior and are of interest as drug delivery vessels, might also influence the structure of cytoskeletal proteins.

We have focused on condensates spreading along actin filaments that have already been bundled by condensates. For the purposes of this study, we have treated the bundle as a uniform cylinder with a radius of the bundle. On planar surfaces, it is known that the texture and microstructural details of the surface influence drop wetting and spreading (42–45). In our system, the bundle is made of filaments that are packed together with condensate creating a surface that is complex and could be porous without any control of the interfilament spacing. Future research will investigate condensate spreading on bundles with different interfilament spacing, expanding our understanding of how drops spread on porous fibers.

## Experimental Methods

### Protein Preparation

Unlabeled monomeric actin (G-actin) is rehydrated from rabbit skeletal muscle lyophilized powder. The powder is briefly centrifuged to ensure all powder is at the bottom of the tube prior to rehydration. The protein is hydrated to 10 mg/mL by adding water and pipette mixing over ice for 15 minutes. Then, the actin solution is dialyzed against the actin storage buffer (in 2 mM Tris, 0.1 mM CaCl_2_, 0.5 mM DTT, 0.2 mM ATP, 1 mM NaN_3_, pH 8) for at least three hours, while spinning on ice. The solution is further dialyzed overnight without stirring at 4°C.

Fluorescently labeled actin is prepared by purifying actin from rabbit skeletal muscle acetone powder purchased from Pel-Freez Biologicals using a protocol adapted from (46). The actin was labeled with tetramethylrhodamine-6-maleimide (TMR). FUS GFP was received as a gift from Avinash Patel and Tony Hyman (30, 47).

Concentration of protein stocks are determined through absorbance measurements using an extinction coefficient for actin of 42680 M^-1^cm^-1^ at 280 nm. The final protein solutions are aliquoted and then flash-frozen in liquid nitrogen and stored at -80°C. Actin is used within three days of thawing, and FUS is used within one day of thawing. After thawing, the proteins are kept at 4°C until needed.

### Experimental Assay

The experimental chamber consists of a borosilicate culture epoxied to the center of a glass coverslip. Prior to epoxying the cylinder, the glass is rinsed twice with ethanol and water. The glass is dried completely with filtered air, and the cylinder is epoxied onto the coverslip. Immediately before adding the sample to the experimental chamber, 5 μL of oil-surfactant solution is added to passivate the coverslip surface and excess surfactant solution is removed via pipette.

To polymerize the actin filaments, actin monomer is added to a buffer solution such that the final composition is 2.64 μM actin monomer (0.264 μM labeled with TMR) in 2 mM Tris-HCl, 2 mM MgCl_2_, 0.50 wt% *β*-mercaptoethanol, 0.3 mM ATP, and 0.40 wt% methylcellulose. The actin is then incubated for at least 20 minutes to allow polymerization into filaments. Then the actin solution is added to the sample chamber. Finally, FUS is added directly to the chamber, such that the final concentration is between 2.64 to 5.28 μM, gently pipette mixed.

### Microscopy

Samples are imaged using a Nikon Eclipse TI2 microscope equipped with a W1 spinning disk confocal head (Yokogawa) via an ORCA-Fusion CMOS camera (Hamamatsu) and a 60×, 1.49 numerical aperture Apo TIRF objective (Nikon). The samples are illuminated using 488 nm and 561 nm laser lines.

### Image Analysis

To determine whether actin and FUS condensates are colocalized in a bundle, we measured the fluorescence intensity associated with each protein along a line that is perpendicular to the long axis of the bundle. To quantify the aspect ratio of condensates, we use ImageJ to measure the lengths of the major axis and perpendicular minor axis of the condensate (48). The aspect ratio is the major axis to minor axis. To determine bundle motion during network coarsening, we used the maximum intensity projection function of ImageJ to compress intensity information from a stack of images into a single image.

### Bundle radius measurement

The bundle radius was measured by first measuring the intensity along a line that is perpendicular to the bundle length and extends at least 20 pixels beyond the bundle on either side. The plot profile function on ImageJ to obtain the intensity across the bundle and length in microns. This information was then plotted and fit to a Gaussian using the following functional form

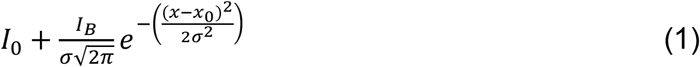

where *I*_0_ is the intensity of the background, *I*_*B*_ is the intensity associated with the bundle, *σ* is the standard deviation of the bundle intensities, and *x*_*0*_ is the location of the maximum intensity. The background intensity is due to the camera and sample dark level that was measured at the tail of the Gaussian. We define the radius of the bundle to the full width at half the maximum of the Gaussian, 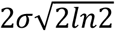. Gaussian fits had an average R^2^ value of 0.992, with a minimum of 0.967. For a single bundle, the radius was measured at 3 points and averaged for 30 bundles.

### Contact angle and shape analysis

The contact angle with the substrate was measured for FUS wrapping around the fiber-like bundle. The formation of a static axisymmetric meniscus with barrel-shaped morphology (Fig. S1) was used as an indication for reaching an equilibrium condition. The droplet shape profile was extracted using MATLAB image processing and edge detection algorithms. The contour was split in four sections – along the long axis at the middle of the fiber, and perpendicular to the fiber at the widest part of the barrel shaped meniscus (Fig. S1). Contour data was separated from the fiber contour at the contact line, and the droplet profile was fitted with the unduloidal function (Eq. 3) based on Young-Laplace equation (Eq. 2):

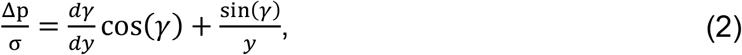

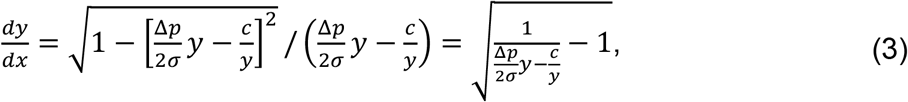

where Δ*p* is the pressure difference between inside and outside of the droplet, *σ* is the interfacial tension, *γ* is the angle that the normal vector to the droplet surface creates with the fiber long axis, *c* is the fitting constant, and 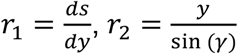 are the principal radii of curvature of the liquid interface. The MATLAB function ode45 was used to find the best fit using the least square method with initial condition at the contact line with the fiber *y*_0_ =*R*_*r*_ and *x*_0_= 0. The fitting parameters 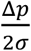 and *c* were defined through measurable parameters as:

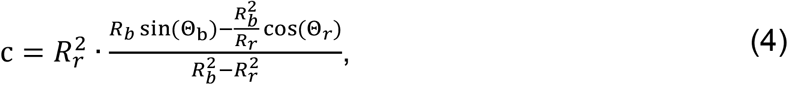

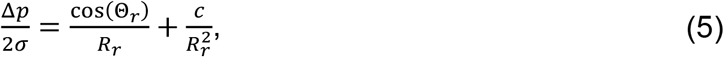

where *R*_*r*_ is the radius of the fiber, *R*_*b*_ is the radius of the barrel-shaped droplet at the cut-off, *θ*_*r*_ is the receding contact angle, *θ*_*b*_ is the angle of the contour against the y axis at the cut-off as shown in Fig. S1 (49).

## Acknowlegements

We thank Avinash Patel and Tony Hyman for the gift of FUS-GFP. This research was supported in part by Clemson Creative Inquiry + Undergraduate Research Program.

## Notes

### Competing Interest Statement

The authors have declared no competing interest.

