## Supplemental Figures for "Capillary-induced bundling of biopolymer networks via protein condensates"

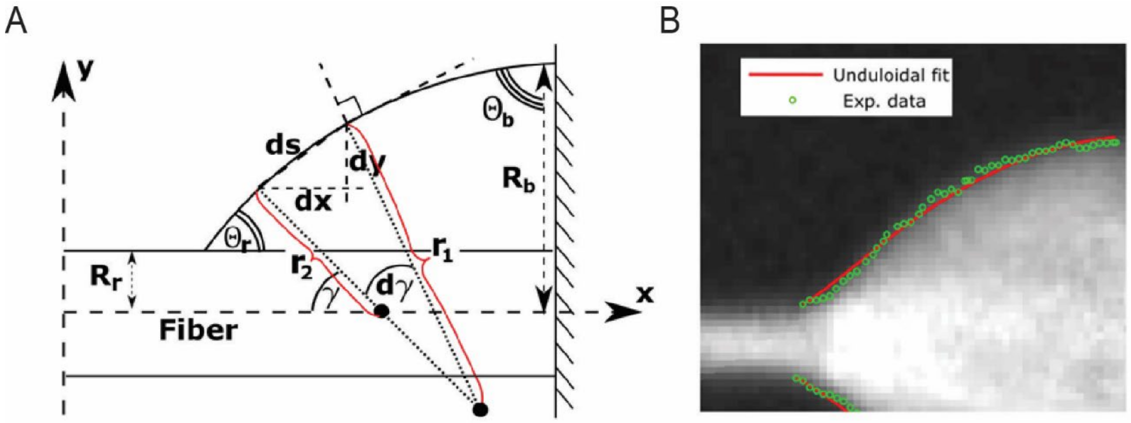


**Fig. S1.** (A) Schematic of static axisymmetric droplet in equilibrium wrapping around fiber with radius $R_r$. The $r_1 = \frac{ds}{dy}$, $r_2 = \frac{y}{\sin(\gamma)}$ are the principal radii of curvature of the liquid interface, $R_b$ – radius of the droplet at the cut-off with angle $\theta_b$ between droplet contour and cut-off, $\gamma$ – the angle between the long axis of the fiber and the normal vector, $ds$ is the differential of arclength and $dx$, $dy$, are differentials of its projectioin on the x and y axes. , $\theta_r$ – receding contact angle that the drop makes with the fiber. (B) Example of droplet contour fit with unduloidal function (Eq. 3) with $R_r = 1.2\ \mu m$, $R_b = 6.5\ \mu m$, $\theta_b = 77\ deg$, and $\theta_r = 31\ deg$.

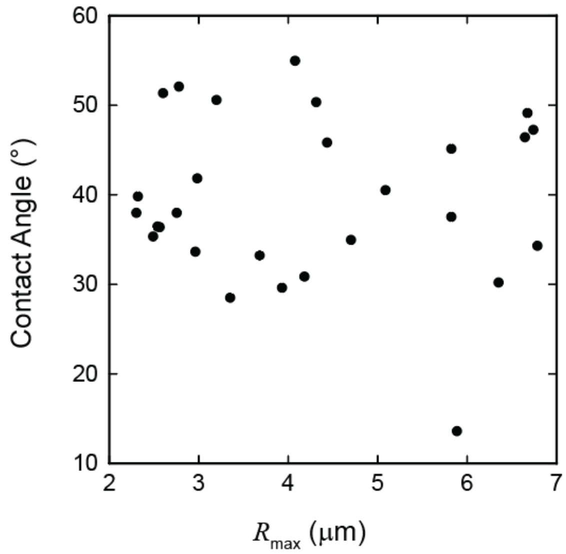


**Fig. S2.** Dependence of the contact angle on the droplet radius, $R_{max}$, calculated for 28 condensates.

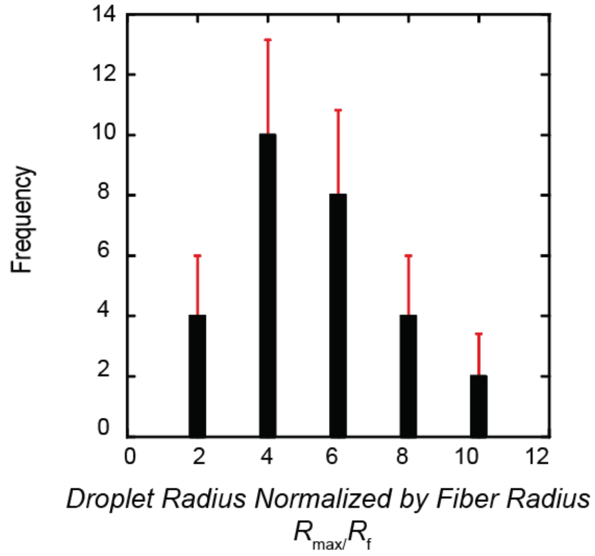


**Fig. S3.** Histogram of the droplet radius normalized by the fiber radius, $\frac{R_{max}}{R_f}$, for 28 condensates. Error is calculated by the standard deviation of the frequency per bin.

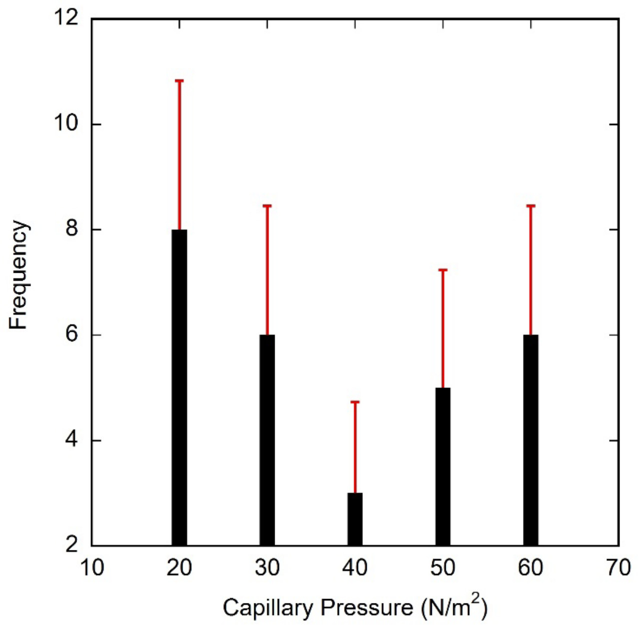


**Fig. S4.** Histogram of the capillary pressure for 28 FUS condensates. Error is calculated by the standard deviation of the frequency per bin.

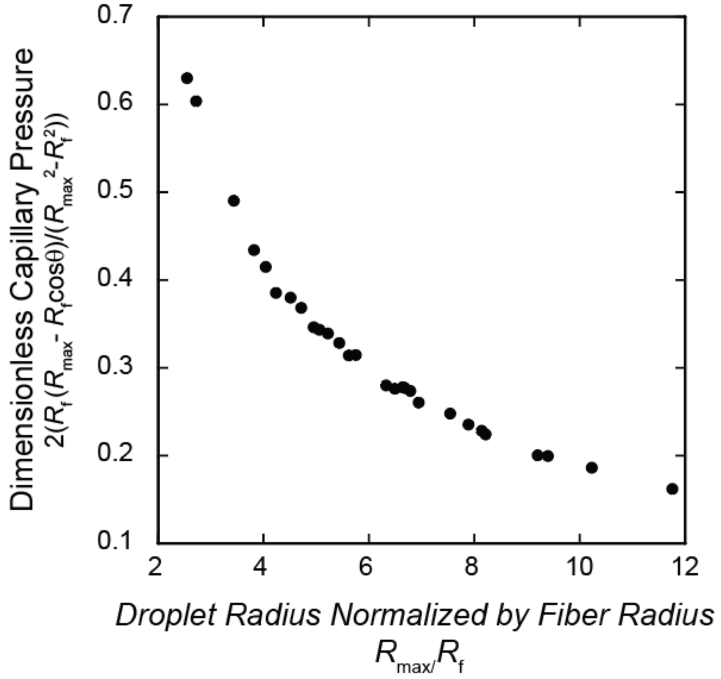


**Fig. S5.** Dependence of the dimensionless capillary pressure, $2\frac{R_f(R_{max}-R_f\cos\theta)}{R_{max}^2-R_f^2}$, on the droplet radius normalized by the fiber radius, $\frac{R_{max}}{R_f}$, calculated for 28 FUS condensates.
